# Comparing fluorometric methods (in vivo vs extracted) for cyanoHAB monitoring in six Rhode Island ponds

**DOI:** 10.1101/2025.06.17.660268

**Authors:** Stephen D. Shivers, Jeffrey W. Hollister, Sophie Fournier, Jakob A. Stankoski, Betty J. Kreakie

## Abstract

Harmful algal blooms caused by cyanobacteria (cyanoHABs) are detrimental to human and environmental health and can be difficult to monitor without specialized training and equipment. A variety of instruments have been developed to measure cyanoHAB indicators (i.e., chlorophyll *a* or phycocyanin) that do not require advanced laboratory processes (e.g., pigment extraction). We compared measurements from five in vivo fluorometers (Turner Trilogy in-vivo module, Turner Fluorosense, Turner Cyanofluor, bbe AlgaeTorch, and bbe Phycoprobe) to results from solvent-based extractions for chlorophyll *a* and phycocyanin at six different waterbodies in Rhode Island. We found a strong relationship between extracted phycocyanin and in vivo fluorometers (R^2^ ranging from 0.78-0.96). We found less consistency between in vivo measurements of chlorophyll *a* and the extracted results (R^2^ between 0.34 and 0.82). Some variability in the chlorophyll *a* results can be explained by differences in the phytoplankton community across the different sampling sites. Phycocyanin results from in vivo fluorometry were also strongly related with cell counts, which implies that phycocyanin measurements from these instruments can be a good proxy for cell counts. Many federal, state, and local entities use cell counts of cyanobacteria to determine when to issue health or contact advisories for waterbodies. Producing accurate cell counts requires highly specialized training/equipment, processing time, and counts can vary greatly between technicians. The results from this study encourage further adoption of in vivo fluorometry and phycocyanin for cyanoHAB monitoring efforts.

## Introduction

Harmful algal blooms caused by cyanobacteria (cyanoHABs) are likely to increase in a warming world with greater inputs of nutrients by humans [1,2] These blooms may impact both human and environmental health, particularly if toxins are being produced. CyanoHAB blooms are often transient in nature, can be difficult to accurately quantify, and occur in diverse waterbodies across large spatial scales [3]. Despite occuring on large scales, effects of blooms have primarily local impacts and can differ in species composition and toxicity based on local conditions [4,5]). As large, centrally managed monitoring efforts are expensive and time consuming to operate, a substantial portion of cyanoHAB monitoring occurs on a smaller, more local level. As neighborhood and lake associations generally do not have the equipment or training to perform extractive analysis of chlorophyll a and/or phycocyanin (pigments found in all algae and specifically cyanobacteria, respectively), alternative measurement equipment has been developed.

A variety of instruments have been designed to measure chlorophyll *a* and/or phycocyanin using in vivo fluorescence without an extraction step. These instruments distinguish between the pigments based on their differing emission spectra. Some instruments (i.e., PhycoProbe) use multiple LEDs at varying wavelengths to cause excitation and emission of different algal type pigments, thus creating a finer scale breakdown of community composition [6]. These instruments can be used in the field, the lab, a combination of both, and have a wide range in cost.

Field instruments may be placed directly in a waterbody or require sample water to be poured into a cuvette and then placed into a handheld fluorometer. Differences in measurement methods represent a potential source of error when comparing instruments. Additionally, instruments may report values in RFUs (raw or relative fluorescence units) or μg/L (concentration) and comparing these values introduces an additional source of error during comparison. In vivo fluorescence can be affected by nonphotochemical quenching causing an underestimation of RFUs as photosystems become saturated with light [7]. Estimation problems can also result from differing phytoplankton/cyanobacteria community composition and growth stage, particularly if colonial cyanobacteria are present as the interior of large colonies will not fluoresce as readily as the exterior [8]. Additionally, the physical properties of the sample water can greatly affect fluorescence readings based on turbidity levels and the presence of colored dissolved organic matter (CDOM). Some instruments offer a yellow substance correction to help account for CDOM while others do not [9]).

Overall, the physical properties and phytoplankton community composition of a waterbody can cause instruments to estimate pigment concentrations differently. How the different instruments handle these various potential situations can affect their accuracy and precision, and the ability to compare across instrument types. The goal of this paper is to compare several handheld and in situ fluorometers with respect to a common benchmark (extracted chlorophyll and phycocyanin) across a variety of waterbodies.

## Methods

### Equipment used for comparison

For this study, we used 5 fluorometers that are commonly used by our collaborators in the northeast US (Table 1). In decreasing order of cost, they are: bbe PhycoProbe, bbe AlgaeTorch, Turner Trilogy (in vivo chlorophyll and orange modules), Turner Cyanofluor, and Turner FluoroSense. Measurements from these fluorometers were compared with the results of a solvent-based extraction for chlorophyll and phycocyanin using a Turner Trilogy with a chla-na and orange module, respectively.

**Table 1:** Manufacturer specifications for the fluorometers that were used in this study.

| Fluorometer | Analysis Method | Measurement Range | Resolution/MDL |
| --- | --- | --- | --- |
| bbe PhycoProbe | cuvette | 0-200 µg chl-a/L | 0.01 µg chl-a/L |
| bbe AlgaeTorch | in situ | 0-500 µg chl-a/L | 0.15 µg chl-a/L |
| Turner Trilogy (in vivo) | cuvette | 0-300 µg/L | 0.025 µg/L |
| Turner Cyanofluor | cuvette | 0-100 µg/L | 0.3 µg/L |
| Turner FluoroSense | in situ | 0-199 µg/L | 1 µg/L |

### Field Sampling Methods

Field samples were collected from 6 ponds in Rhode Island (Figure 1). Samples from Barber, Indian, and Yawgoo were collected on 2021-10-06 and samples from Curran, Mashapaug, and Warwick were collected on 2021-10-22. Surface samples were collected in triplicate by wading into each pond to a minimum depth of 2 feet to avoid collecting near the sediment. If a surface scum was present, the scum was gently brushed aside to avoid being collected in the sample bottles. Samples were collected in acid-washed 1 L amber bottles, placed in a cooler with ice, and stored at 4° C until analysis. At the same location as sample collection, measurements of chlorophyll and phycocyanin were made using the AlgaeTorch and the FluoroSense.

**Fig 1:**
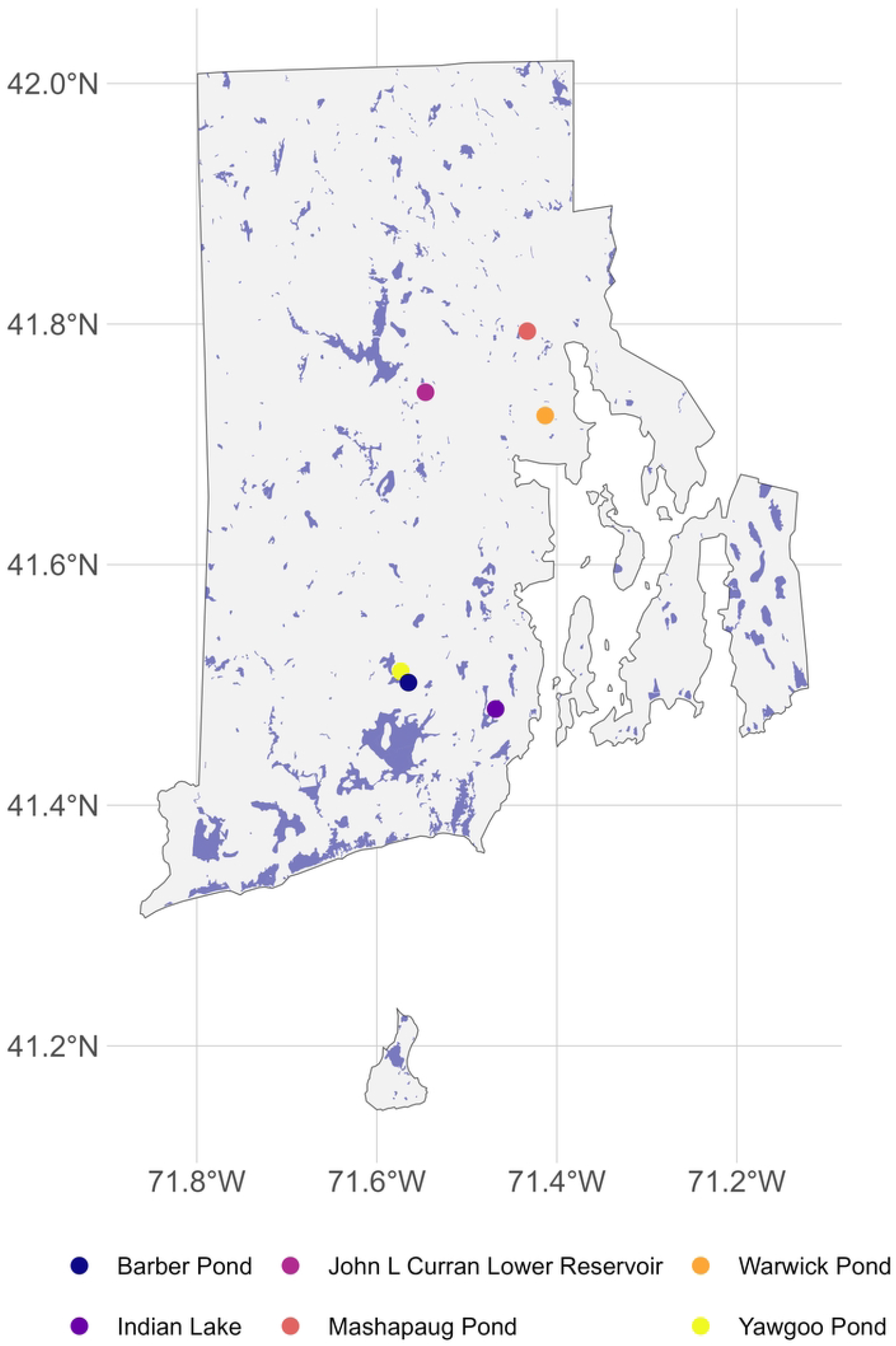
Six field sampling locations in Rhode Island, USA.

### Analysis of field samples

Within 24 hours of collection, field samples were analyzed using the PhycoProbe, CyanoFluor, and the Trilogy (in-vivo), as well as filtered onto 0.7 μm pre-ashed glass fiber filters and frozen at -20° C for solvent-based extraction. Sample water filtered through a 0.22 μm syringe filter was used to correct for yellow substances on the PhycoProbe, CyanoFluor, and Trilogy in vivo module. Chlorophyll *a* was extracted by placing frozen GF/F filters in centrifuge tubes with 90% acetone, then placing the tubes for a 20-minute period in a sonicating water bath to lyse cyanobacterial cells [10]. Phycocyanin was extracted from frozen GF/F filters using a 50 mM phosphate buffer as the extraction solvent and following a 15-minute period in a sonicating water bath to lyse cyanobacterial cells [11]. Extracted samples were analyzed using a Turner Trilogy with a chla na module (chlorophyll) and an orange module (phycocyanin).

### Phytoplankton Counts

All phytoplankton samples were collected on 2021-10-06. Water samples for phytoplankton identification and enumeration were decanted from field collected samples into 250 ml amber HDPE bottles and were preserved with 25% glutaraldehyde. Samples were stored at 4° C and shipped the week after collection on ice to Phycotech, Inc. A minimum of 400 natural units were counted per sample, and quantitative biovolume estimates were also performed for each sample. A full description of methods and a quality assurance plan are available from Phycotech,

Inc. (https://www.phycotech.com/Portals/0/PDFs/PhycoTech_INFO_PACKET.pdf).

### Data Analysis

To compare extracted concentrations and cell counts to the various instruments we used simple linear regression and scatterplots. We derived the coefficient of determination from the regressions as a measure of fit and also qualitatively compared the device measurements to the extracted concentrations and phytoplankton counts with scatterplots. We do not compare slopes nor look for a slope of 1 as we have little expectation that the extracted concentrations would equal the values from the various devices given the difference in units and the different RFU returns across the devices. All data and code for this analysis are available via GitHub at https://github.com/usepa/fluoroproj. This repo has also been archived at Zenodo [12].

## Results

### Chlorophyll

Extracted chlorophyll measurements ranged from a mean of 1.3 μg/L in Indian Lake to a mean of 32.2 μg/L in Mashapaug Pond (Table 2). Chlorophyll measurements for all instruments followed the same general pattern as the extracted chlorophyll values (Figure 2). The Algaetorch and Phycoprobe measurements exhibited a strong relationship with extracted chlorophyll concentrations with R^2^ values of 0.82 and 0.81, respectively. The Cyanofluor (R^2^=0.35) and Trilogy in vivo (R^2^=0.34) measurements increased as extracted chlorophyll concentrations increased, but the relationship was not as strong compared to the other instruments.

**Table 2:** Summary (mean and standard deviation) of extracted chlorophyll and phycocyanin from Turner Trilogy for waterbodies sampled in Rhode Island.

| waterbody | chlorophyll ($\mu\text{g/L}$) | phycocyanin ($\mu\text{g/L}$) |
| --- | --- | --- |
| Barber | $4.5 \pm 0.2$ | $0.2 \pm 0.0$ |
| Curran | $19.6 \pm 0.2$ | $1.5 \pm 0.4$ |
| Indian | $1.3 \pm 0.0$ | $0.2 \pm 0.0$ |
| Mashapaug | $32.2 \pm 3.6$ | $10.5 \pm 2.6$ |
| Warwick | $20.0 \pm 1.0$ | $2.4 \pm 0.5$ |
| Yawgoo | $2.8 \pm 0.2$ | $1.0 \pm 0.1$ |

**Fig 2:**
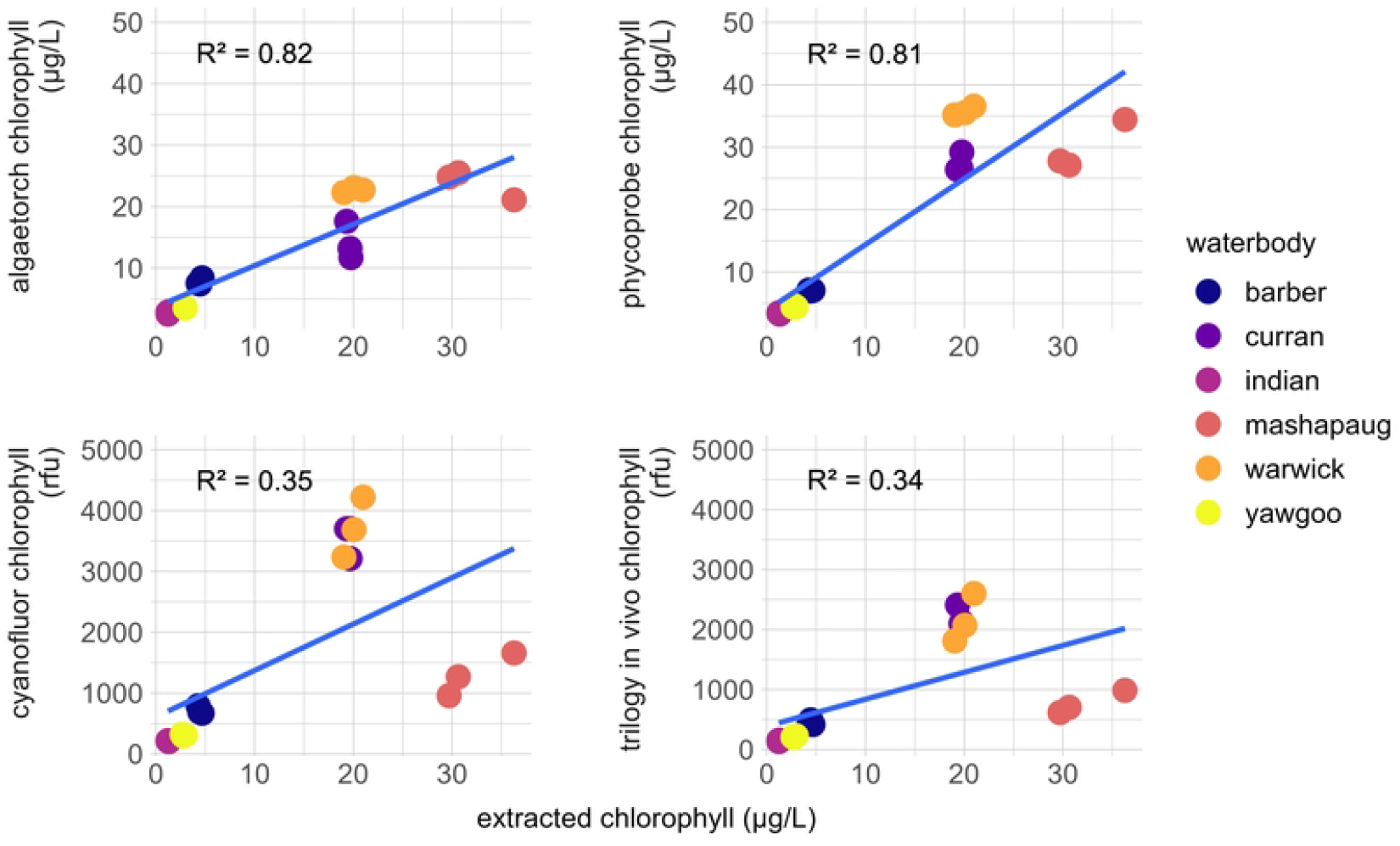
Comparison of fluorometer measurements of chlorophyll by waterbody. Blue lines represent the best fit regression line. R^2^ on plot is the coefficient of determination for this line.

### Phycocyanin

Extracted phycocyanin measurements ranged from a mean of 0.2 μg/L in Indian Lake and Barber Pond to 10.5 μg/L in Mashapaug Pond (Table 2). The Algaetorch (R^2^=0.86) and Phycoprobe (R^2^=0.84) were strongly related to extracted phycocyanin concentrations (Figure 3). Phycocyanin measurements for the Fluorosense (R^2^=0.96) and the Cyanofluor (R^2^=0.78) were also strongly related with extracted phycocyanin.

**Fig 3:**
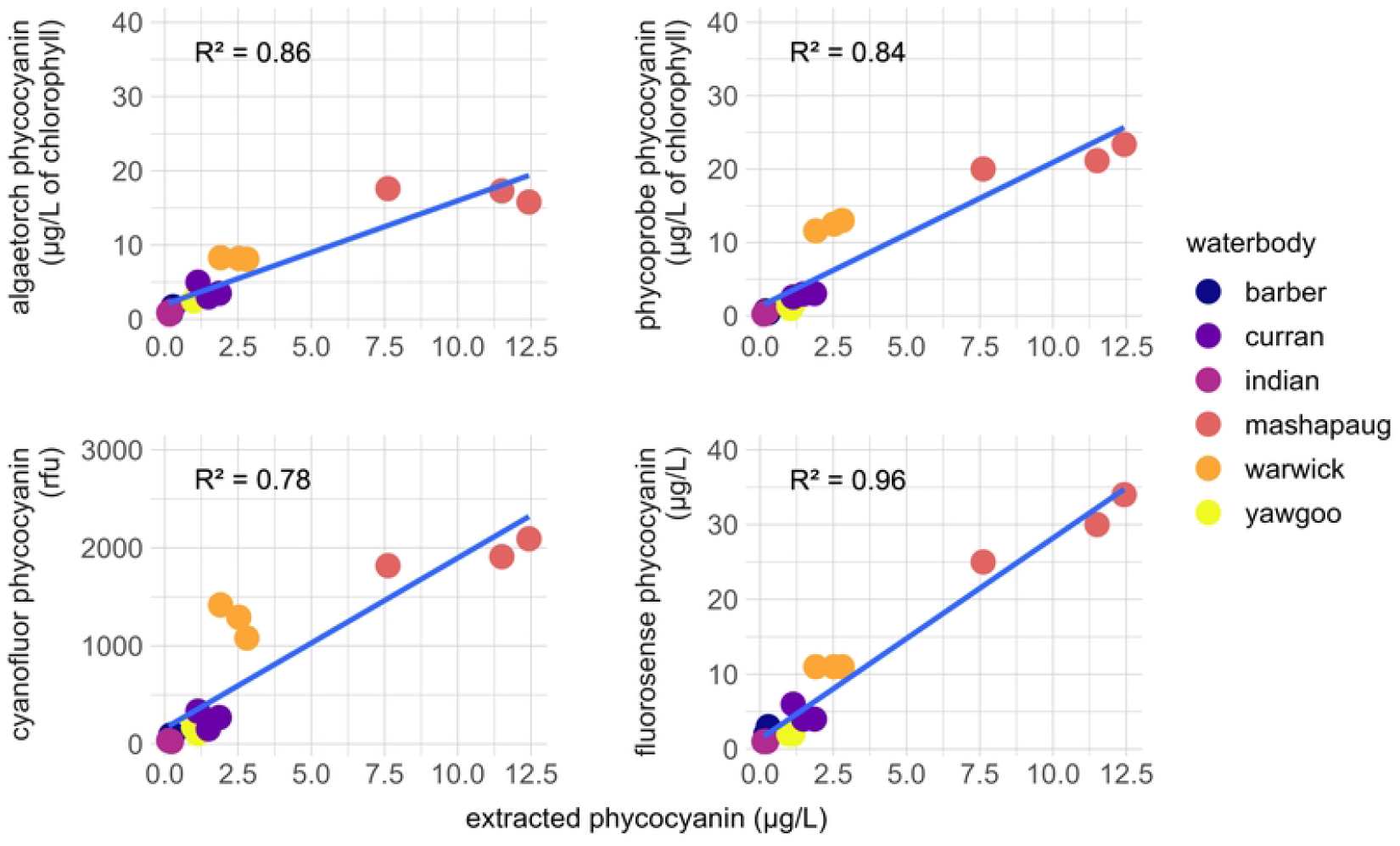
Comparison of fluorometer measurements of phycocyanin by Rhode Island waterbody. Blue lines represent the best fit regression line. R^2^ on plot is the coefficient of determination for this line.

### Phytoplankton Counts

Cyanophytes were the dominant taxa at all waterbodies with the highest concentrations observed at Curran (38,019 natural units/mL), Mashapaug (89,806 natural units/mL), and Warwick (45,847 natural units/mL)(Figure 4). Total concentrations were highest at Mashapaug (92,527 natural units/mL), Warwick (70,728 natural units/mL), and Curran (40,428 natural units/mL).

**Fig 4:**
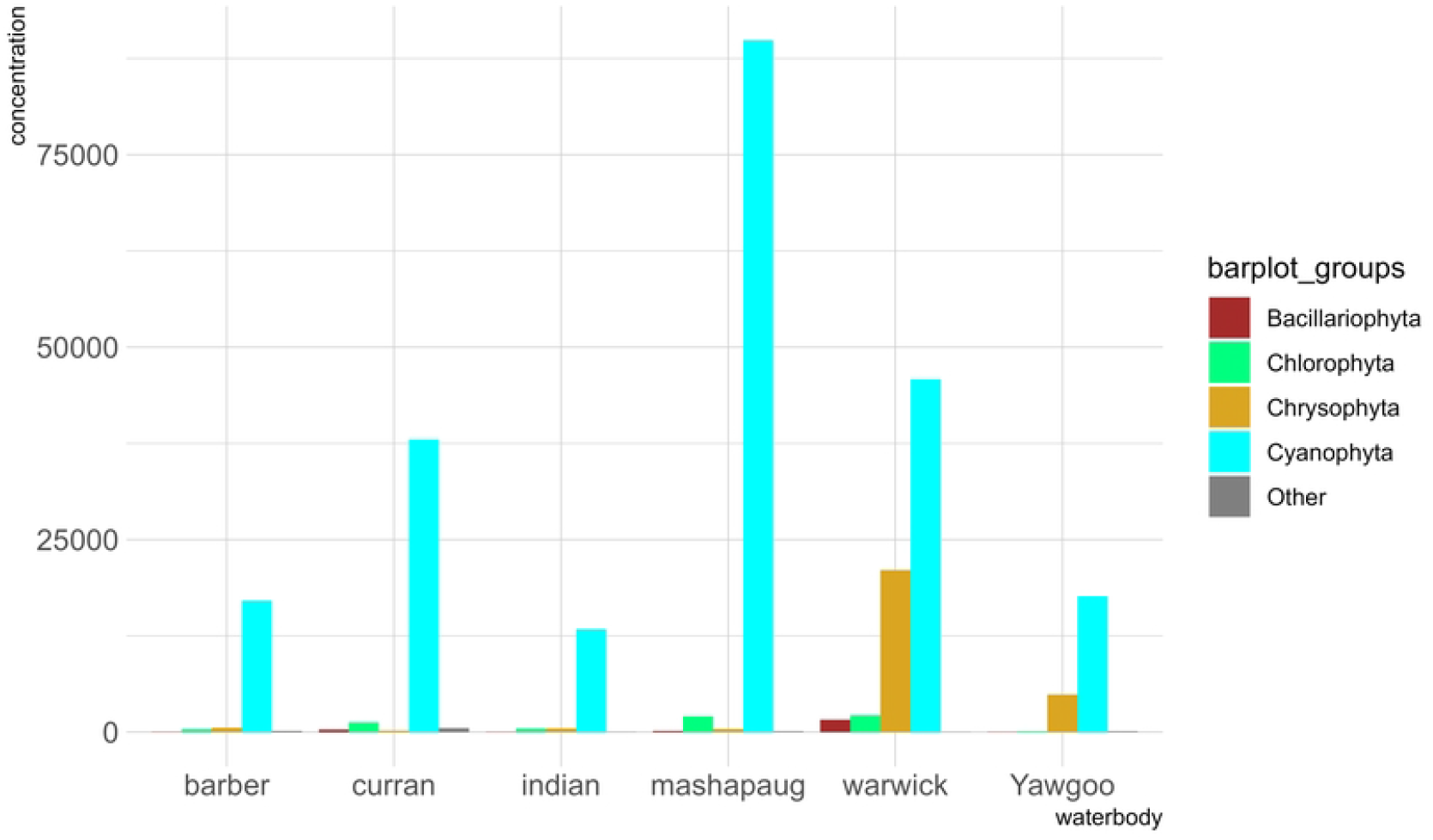
Cell counts (cells/mL) of phytoplankton samples from six waterbodies in Rhode Island.

Extracted chlorophyll *a* exhibited the strongest relationship with cyanobacterial cell counts (R^2^=0.92) followed by the Algaetorch (R^2^=0.78) and the Phycoprobe (R^2^=0.59). The Cyanofluor (R^2^=0.12) and Trilogy in vivo module (0.11) exhibited a weaker relationship (Figure 5). Phycocyanin measurements from all instruments were strongly related to cell counts (R^2^ between 0.9 and 0.97)(Figure 6).

**Fig 5:**
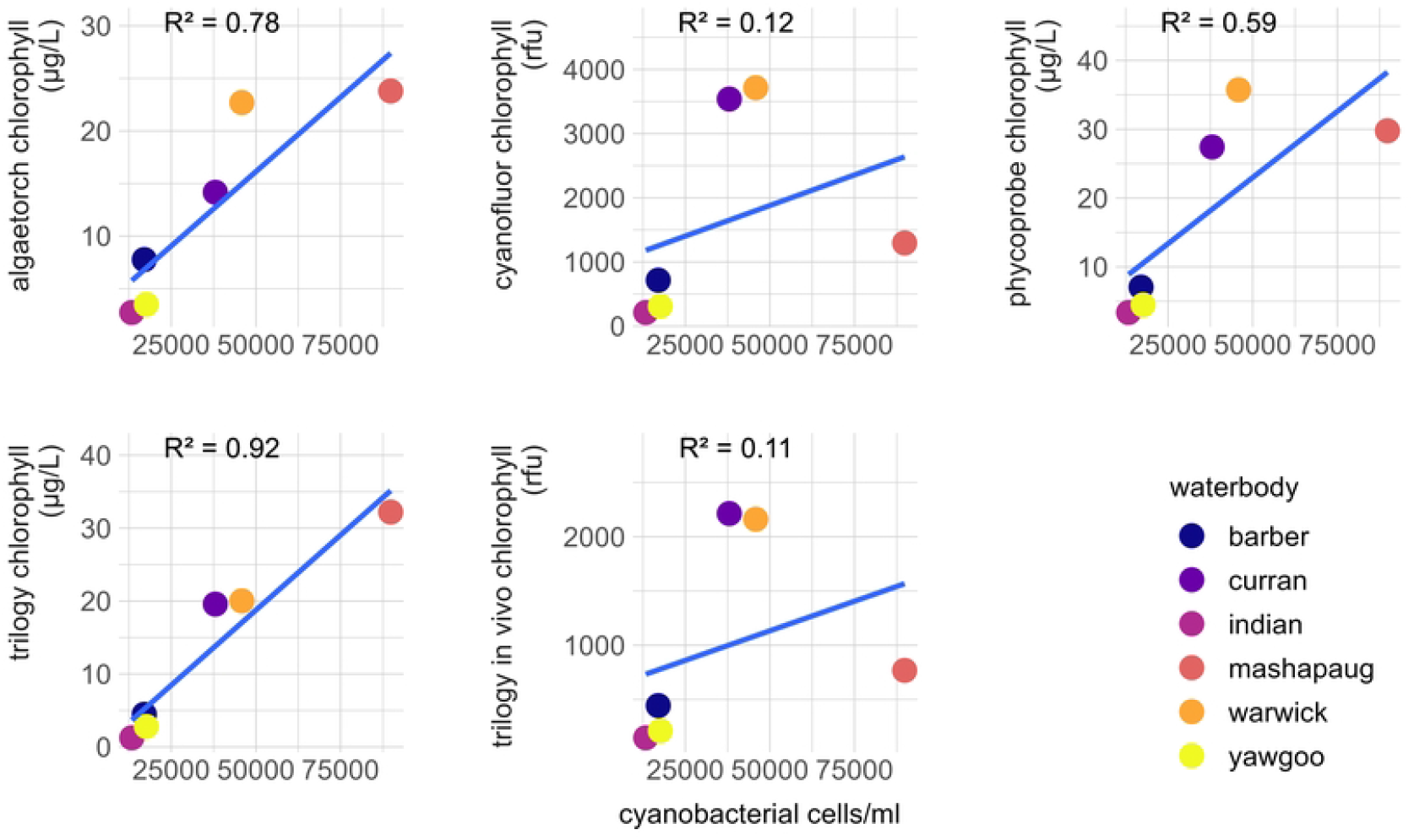
Comparison of fluorometer chlorophyll to cyanobacteria cell counts. Blue lines represent the best fit regression line. R^2^ on plot is the coefficient of determination for this line.

**Fig 6:**
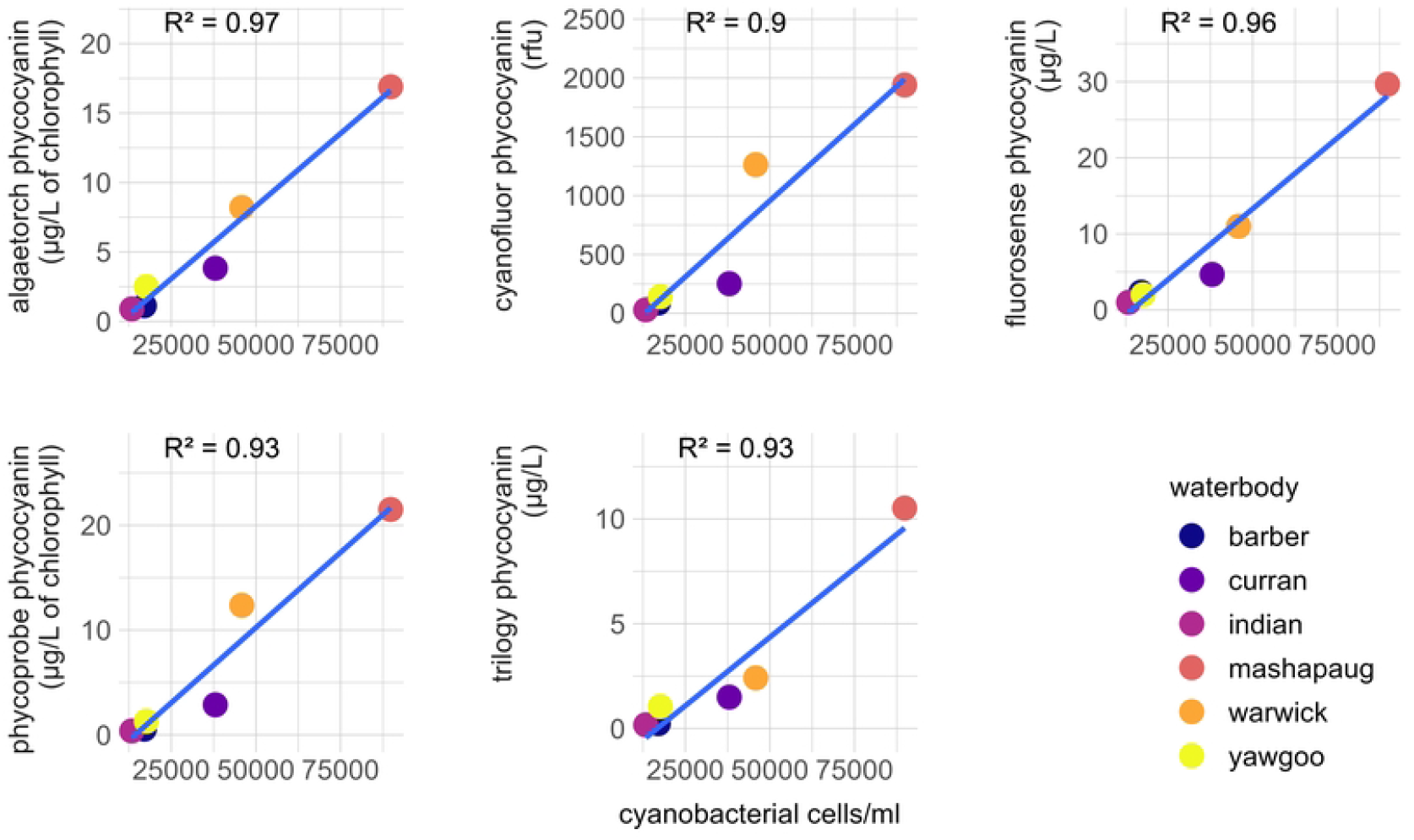
Comparison of fluorometer phycocyanin to cyanobacteria cell counts. Blue lines represent the best fit regression line. R^2^ on plot is the coefficient of determination for this line.

## Discussion

All fluorometers performed well when comparing phycocyanin measurements to extracted measurements, and these results agree with previous studies that supported the use of phycocyanin as an indicator of HABs caused by cyanobacteria[13–15].The Fluorosense had the strongest relationship between the waterbodies; however, the Fluorosense overestimated the actual concentrations as compared to extracted values (0-35 μg/L vs 0-12.5 μg/L). Hambdhani et al. (2021) also found that the Fluorosense performed well but overestimated chlorophyll *a* at higher algal concentrations [16]. Therefore, using this particular instrument to enforce set criteria across a variety of waterbodies with no prior knowledge of expected values would probably not be recommended without further research.

The Algaetorch and Phycoprobe yielded similar measurements and also overestimated phycocyanin concentrations compared to extracted concentrations; however, the output from these instruments is not a direct measure of phycocyanin. Instead, these instruments measure the percentage of the chlorophyll that is derived from cyanobacteria, which makes a comparison with other instruments that directly measure phycocyanin difficult. The Cyanofluor output was in RFUs and was not directly comparable to extracted phycocyanin without creating a standard curve for the instrument using known phycocyanin standard concentrations, which was beyond the scope of this study. A study by Thomson-Laing et al. (2020) did convert RFUs to μg/L and found strong correlation between cyanobacteria biovolume and phycocyanin concentration for eleven cyanobacteria cultures [17]. They also found stronger relationships for environmental samples in lakes that were cyanobacteria dominated compared to more diverse lakes. All of these instruments could be used to effectively monitor for the presence of cyanobacteria by measuring more direct parameters of cyanobacteria than chlorophyll *a*.

All of the fluorometers exhibited a positive relationship with chlorophyll, but the Algaetorch and Phycoprobe had a much stronger relationship. Weaker relationships by the Cyanofluor and Trilogy in vivo module was driven by the underestimation of chlorophyll in Mashapaug Pond and possible overestimation in Warwick and Curran. Phycocyanin concentrations and cell counts of cyanobacteria were the highest in Mashapaug and it is likely that these high concentrations were causing underestimation of the in vivo concentrations of chlorophyll [18]. Therefore, the Algaetorch and Phycoprobe are more successful in properly estimating chlorophyll across a range of different waterbodies, but they are also the most expensive instruments. A study by Silva et al. (2016) found that the Fluoroprobe (a similar instrument to the Phycoprobe) was correlated with cell counts at chlorophyll *a* concentrations below 100 μg/L but not above, but the present study did not have study ponds that exceed 100 μg/L [19].

This study supports the use of commercially available fluorometers for monitoring cyanoHABs when other techniques are not feasible. In lieu of extractive techniques, in vivo chlorophyll *a* is typically used for routine monitoring of cyanoHABs, but this study found in vivo phycocyanin to be better related to extracted phycocyanin than the relationship between in vivo and extracted chlorophyll *a* across a variety of instruments. Therefore, measuring in vivo phycocyanin may be a better solution for HAB monitoring. There are a variety of instruments capable of measuring phycocyanin using in vivo fluorometry across a spectrum of prices. We hope this study will encourage further adoption of using in vivo phycocyanin as alternative to cell counts and in situations where extractive techniques are not able to be used.

## Acknowledgments

We are grateful to Hilary Snook, Darryl Keith, Steve Rego, Kara Godineaux, Marty Chintala, and Tim Gleason for constructive reviews of this paper. The views expressed in this article are those of the authors and do not necessarily represent the views or policies of the U.S. Environmental Protection Agency. Any mention of trade names, products, or services does not imply an endorsement by the U.S. Government or the U.S. Environmental Protection Agency. The EPA does not endorse any commercial products, services, or enterprises. This contribution is identified by the tracking number ORD-060125 of the Atlantic Coastal Environmental Sciences Division, Office of Research and Development, Center for Environmental Measurement and Modeling, US Environmental Protection Agency. Lastly, special thanks to Gracie, Anastasia, and Lilac for cheer leading and support during challenging analysis times.

